# ADAR1 can drive Multiple Myeloma progression by acting both as an RNA editor of specific transcripts and as a DNA mutator of their cognate genes

**DOI:** 10.1101/2020.02.11.943845

**Authors:** Rafail Nikolaos Tasakis, Alessandro Laganà, Dimitra Stamkopoulou, David T. Melnekoff, Pavithra Nedumaran, Violetta Leshchenko, Riccardo Pecori, Samir Parekh, F. Nina Papavasiliou

## Abstract

RNA editing is an epitranscriptomic modification of emerging relevance to disease development and manifestations. ADAR1, which resides on human chromosome 1q21, is an RNA editor whose over-expression, either by interferon (IFN) induction or through gene amplification, is associated with increased editing and poor outcomes in Multiple Myeloma (MM). Here we explored the role of ADAR1 in the context of MM progression, by focusing on a group of 23 patients in the MMRF CoMMpass Study for which RNAseq and WES datasets exist for matched pre-and post-relapse samples. Our analysis reveals an acquisition of new DNA mutations on disease progression at specific loci surrounding the sites of ADAR associated (A-to-I) RNA editing. These analyses suggest that the RNA editing enzyme ADAR1 can function as a DNA mutator during Multiple Myeloma (MM) progression, and further imply that guide-targeted RNA editing has the capacity to generate specific mutational signatures at predetermined locations. This dual role of RNA editor and DNA mutator might be shared by other deaminases, such as APOBECs, so that DNA mutation might be the result of collateral damage on the genome by an editing enzyme whose primary job is to re-code the cognate transcript toward specific functional outcomes.

## INTRODUCTION

The **A**denosine **D**eaminase **A**cting on **R**NA (ADAR) family consists of two enzymes (ADAR1 and −2) with demonstrated deaminase activity and one (ADAR3) for which catalytic deamination activity has not been demonstrated, but which might have other functions (Valente and Nishikura, 2005; Gallo et al., 2017; Oakes et al., 2017; Mladenova et al., 2018). The focus of this study, ADAR1, is ubiquitously expressed. ADAR1 deaminates Adenosine to Inosine (A-to-I), which is recognized by cellular machineries as a Guanosine (G). This molecular phenomenon is called RNA editing or A-to-I editing, and it is widespread in the transcriptome.

ADAR1-mediated RNA editing is required for survival: even transient loss of the enzyme in mouse leads to organism-wide toxicity and death (Pestal et al., 2015). This is in part because the editing enzyme recognizes and targets “self” dsRNAs (Athanasiadis, Rich and Maas, 2004) which upon editing, are no longer detected by cellular nucleic acid sensors as “foreign”. Loss of ADAR1 therefore leads to improper recognition of “self” dsRNA, which triggers an interferon response, and cell death in most contexts.

ADAR1 is highly overexpressed in all human cancers (in comparison to cognate healthy tissue); kidney chromophore tumors are an exception to that rule (Han et al., 2015; Xu and Öhman, 2019). Thus, the vast majority of human tumors are highly edited, especially at inverted *Alu* repeats (which provide the necessary dsRNA target structure): indeed, editing at *Alus* has been used to generate an “index” (Alu Editing Index or AEI) that can be used comparatively to stage tumors (Bazak, Levanon and Eisenberg, 2014; Paz-Yaacov et al., 2015; Roth, Levanon and Eisenberg, 2019). Recently, CRISPR/Cas screens revealed that tumors are vulnerable to ADAR1 loss, which coincides with a substantial interferon response, and with tumor cell death (Gannon et al., 2018). But even in tumors where the IFN pathway is abrogated, loss of ADAR1 is still consequential, possibly because it reduces the informational heterogeneity of a tumor (Paz-Yaacov et al., 2015; Pecori et al, submitted). These recent experiments have motivated a search for ADAR1 inhibitors as cancer therapeutics.

Here, we focus on Multiple Myeloma (MM), a malignant cancer of plasma cells (Laganà et al., 2018). One of the common features of MM (occurring in roughly 40% of newly diagnosed patients) is the amplification of chromosome 1q21, which is associated with poor prognosis (Klein et al., 2010; Nemec et al., 2010; Chang et al., 2011). The 1q21 locus is also the genomic location of ADAR1, and when this chromosomal fraction is amplified in MM, ADAR1 expression is increased, as is editing (Lazzari et al., 2017; Teoh et al., 2018). Because of this link between 1q21 amplification and prognosis in MM, MM has been one of the tumors in which ADAR1 editing has been relatively well studied (Lazzari et al., 2017; Teoh et al., 2018). A similar mechanism of ADAR1 upregulation by 1q amplification has also been described in breast cancer (Fumagalli et al., 2015). Here we turn our attention to the role of ADAR1 in MM disease progression, and describe a new role for ADAR1, not only as an RNA editor but also as a DNA mutator.

## RESULTS

### ADAR1 expression in Multiple Myeloma is driven either by 1q amplification or by interferon signaling

The amplification of 1q identifies a subclass of MM patients characterized by a more aggressive disease course and poor prognosis. The ADAR1 gene resides in 1q21 and previous studies have reported higher expression of ADAR1 in patients with 1q amplification and have shown worse progression-free (PFS) and overall survival (OS) in patients with high ADAR1 expression (Lazzari et al, 2017; Teoh et al, 2018). We analyzed Whole-Genome Sequencing (WGS), Whole-Exome Sequencing (WES) and RNA-Seq data from 526 newly diagnosed MM patients in the MMRF CoMMpass study (https://research.themmrf.org/). Our analysis confirmed worse PFS and OS for patients with 1q copy number (CN) gain (>2 copies) (Fig. 1A and Fig. S1A, CN>2 shown as ADAR+) and that ADAR1 expression was significantly higher for the ADAR+ patients (Wilcoxon Rank Sum Test p < 10^−3^) (Fig. 1B). However, we also observed several patients with high expression of ADAR1 and no ADAR1 copy number gain (Fig. 1C). Differential expression analysis revealed a significant enrichment of interferon-related pathways (IFN) in this group of patients (adj-p < 10^−7^). These findings strongly suggest that ADAR1 overexpression is a key driver in MM, and this can be achieved either through 1q21 amplification or through IFN induction (and subsequent upregulation of the ADAR1 p150 isoform, which is an interferon-stimulated gene (ISG) (George, Wagner and Samuel, 2005). A similar situation has been reported in breast cancer as well (Fumagalli et al., 2015).

**Figure 1.**
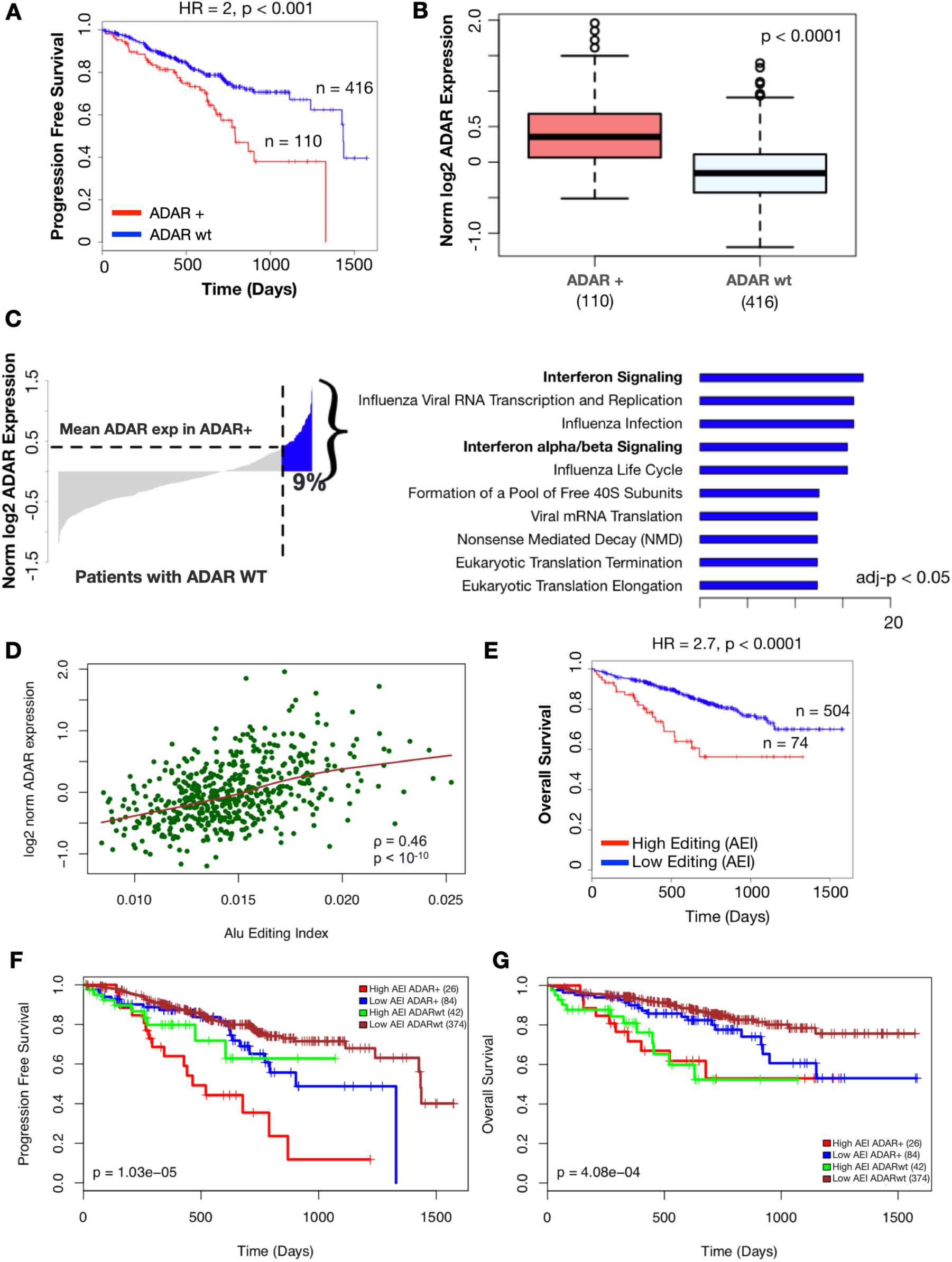
ADAR1 expression and ADAR1-mediated editing are significantly associated with poor prognosis in Multiple Myeloma. (A) Progression-free survival for patients with ADAR1 copy number (CN) gain (>2) (ADAR+, n=110) and ADAR1 Wild Type patients (ADARwt, n=416), (B) ADAR1 expression between patients with ADAR1 CN gain and ADARwt patients (normalized log2 counts), (C) Interferon signaling pathways are enriched in ADARwt patients with ADAR expression above the mean of patients with ADAR1 CN gain (ADAR+; 9% of the patients – also observed as outliers in ADARwt, Fig. 1B), (D) Alu Editing Index (AEI) is significantly correlated (p<10^−10^) with ADAR1 expression, (E) Overall survival analysis for High vs Low AEI patients, (F) Progression-free survival and (G) overall survival for patients grouped by AEI and 1q status: AEI 1q21+ (red), High AEI 1q21WT (green), Low AEI 1q21+ (blue), Low AEI 1q21WT (deep red).

### Elevated ADAR1-mediated RNA editing (quantified with the Alu Editing Index or AEI) is strongly associated with poor patient survival

To quantify ADAR1-mediated A-to-I RNA editing in a sample-wise fashion, we calculated Alu Editing Index (AEI) for an extended set of 590 patients in the MMRF CoMMpass study. Our results showed that AEI is strongly correlated with ADAR1 global expression across the patient cohort (Fig. 1D) and that patients with high AEI had significantly lower PFS and OS (Fig. S1B, Fig. 1E). ADAR2 is expressed at low levels in this cohort. Furthermore, AEI could significantly discriminate between good and poor prognosis even within the group of patients with 1q gain (PFS: p=1.03×10^−5^, OS: p=4.08×10^−4^), demonstrating that high RNA editing, as measured by AEI, can be an adverse prognostic factor independently of such alterations (Fig. 1F, 1G). Concordantly with the analysis of ADAR expression, pathway enrichment analysis revealed that patients with high AEI were characterized by up-regulation of interferon signaling, supporting the contribution of pro-inflammatory signaling to ADAR expression and activity, and by down-regulation of immunoglobulin genes and pathways that are relevant in MM, such as the unfolded protein response (UPR) and glycosylation (Fig. S1C).

### A dual role for ADAR1 as RNA editor and DNA mutator in Multiple Myeloma

We had previously noted that, within a cohort of patients at a specific disease stage, the list of known driver mutations at the population level was broadly similar with lists of ADAR edited sites per patient (this was true for DLBCL – Pecori et al, submitted; and similarly, for MM (Figure S2). We therefore wondered whether ADAR1 was capable of not only editing RNA but also mutating DNA. Fundamentally this would be possible as the ubiquitous ADAR1 isoform p110 contains Z-DNA binding domains (Zinshteyn and Nishikura, 2009) that enable ADAR1 to bind to genomic DNA during transcription (Herbert et al., 1998; Schwartz et al., 1999; Herbert 2019). Additionally, ADAR1 was recently shown to be capable of mutating DNA within DNA:RNA hybrids (R-loops; Zheng, Lorenzo and Beal, 2017). We therefore asked whether we could identify ADAR-mediated mutations, using principles derived from mechanistic knowledge of how ADAR1 could target DNA (**Fig. 2A**): specifically, we hypothesized that any ADAR1 editing on DNA would happen within R-loops where the RNA itself would be a canonical target of the ADAR1, so that A-to-I editing on the transcript could also generate A-to-I (hypoxanthine) mutations in the transcribed (-) DNA strand, which we would then identify as a T-to-C mutation on the reference (+) strand (or as a T-to-N mutation, in case DNA repair has failed to properly correct the hypoxanthine on the negative strand).

**Figure 2.**
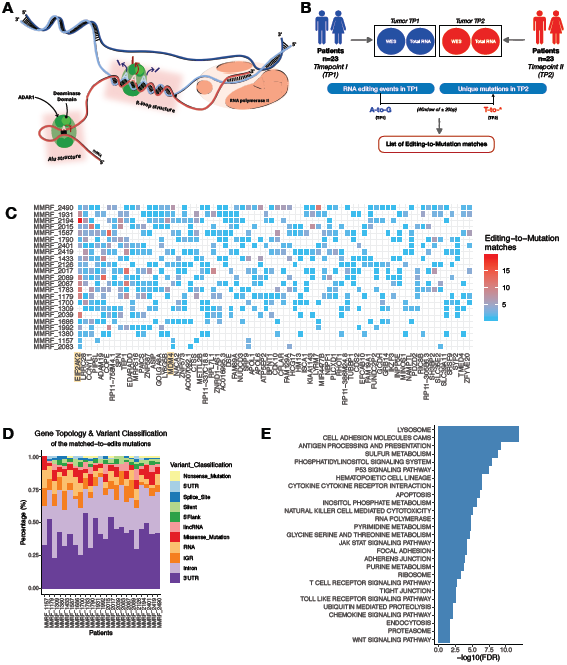
ADAR1 can function as DNA mutator in Multiple Myeloma. (A) ADAR1 edits double-stranded RNA co-transcriptionally mostly within *Alu* elements. Here, we hypothesize that ADAR1 can access R-loops (hybrids of nascent RNA and the transcribed strand) that occur in the transcription bubble and by targeting the RNA within the loop it can mutate DNA in the vicinity. (B) WES and total RNA-seq data, from two successive timepoints (Timepoint I or TP1 and Timepoint II or TP2) of 23 patients from the CoMMpass MMRF study, were processed to call RNA editing events and gDNA mutations. A-to-G editing events of TP1 were matched to unique T-derived (T-to-*) mutations in TP2, which fall within a window of 20bp up- or down-stream the editing event (*see Methods for details*). (C) Analysis of matching editing events in TP1 to unique mutations in TP2 within conservative windows of ±20bp from the editing event return robust editing-to-mutation matches per patient and per gene. In this heatmap, ADAR1-related mutations in the top-50 genes shared by at least 25% of the patients. The color-scale represents the counts of editing-to-mutation matches. (D) Editing to mutation matches are classified by gene topology per patient. The vast majority occurs in 3’UTRs and introns, characteristic locations of ADAR1-dependent editing. (E) Pathway enrichment analysis for the editing-to-mutation candidates across the 23-patient dataset.

Based on what is known regarding other deaminases (e.g. the APOBEC family), we would expect that mutations would happen at a rate that is substantially lower than the rate of the editing event (for example, APOBEC1 mutates DNA at a rate of 1 in 10,000 bp per population doubling (Saraconi et al., 2014), but can edit cognate RNA at rates approaching 100% for some transcripts). We therefore reasoned that should editing-associated mutational events exist, they would be most prominent in the context of relapse. We thus focused our analysis on 23 patient samples from the COMPASS database, for whom matched genomic (WGS) and transcriptomic (RNAseq) data were available for at least two timepoints (TP1 and TP2, pre- and post-treatment or relapse respectively) (Fig. 2B). We then asked whether we could correlate RNA editing (in transcripts in TP1) with DNA mutation (in genomic DNA, in TP2). Specifically, we focused on positions of A-to-I editing within a transcript at diagnosis, and compared those to instances of A-to-G mutation on the transcribed (-) strand of the cognate gene at relapse (read as T-to-C or more generally, T-to-N mutations on the + strand), within a 41bp window centered on the edited base (we termed these “editing-to-mutation” matches) (Fig. 2B). Within the 23-patient cohort, we ranked the genes with RNA editing shared by at least 25% of the patients according to the number of mutations within the editing windows. Among the top-50 genes in this list, we found a number of cancer-related new targets of RNA editing that include *Mdm4* and *Eif2ak2. Mdm4* encodes a well-known TP53 inhibitor (Danovi et al., 2004). *Eif2ak2* encodes **P**rotein **k**inase regulated by **R**NA or PKR, an interferon stimulated gene (ISG) that plays a central and well-established role in the response against RNA viruses: its activation by dsRNA leads to translational shutdown (Alisi et al., 2005). PKR also plays a crucial role in tumor suppression through interaction with p53 (Yoon, et al., 2010). Both these transcripts have been reported as edited in the RADAR and REDIportal databases in the past (Ramaswami and Li, 2013; Picardi et al., 2016), supporting our finding that they are indeed targets of ADAR1, which we have confirmed experimentally for the *Eif2ak2* transcript in MM cell lines (Figure S3). More importantly, we observed a concurrence between RNA editing (pre-relapse) and the emergence of DNA mutation (post-relapse), with *Eif2ak2 (PKR)* being the top shared target of post-relapse mutation this cohort (Fig. 2C). The DNA mutations we identified are mostly found in 3’UTRs, where ADAR1 typically edits within *Alu* repeats (Fig. 2D), and about 80% of them are found in genomic topologies, known to form R-loops, as defined by publicly available DRIP-seq datasets (Fig. S4). Additional cohort- and target-wise pathway enrichment analysis showed enrichment in signaling pathways associated with apoptosis and tumorigenesis (p53 signaling, JAK-STAT among others) (Fig. 2E), which suggests that ADAR1 might power disease progression by modulating the outcome of these pathways, either by editing or by mutation.

### Disease progression in Multiple Myeloma is facilitated either through elevated RNA editing activity or through ADAR1-associated DNA mutation

Due to the robust concurrence of the editing-to-mutation candidates with key components of tumorigenesis (Fig. 2C,E), we asked whether we could monitor the ADAR1-mediated generation of mutations (in TP2) by evaluating the abundance of editing activity (AEI) in TP1, in the 23-patient cohort. We then plotted editing-to-mutation instances grouped per gene and per patient. We found a strong correlation with the patients’ AEI in TP1 (R=0.66, p<10^−3^), but not in TP2 (Fig.3A). The strong correlation between editing-to-mutation counts and AEI in TP1 is consistent with our hypothesis that ADAR1 might induce DNA off-targets during its canonical RNA editing activity. The observation that there is no correlation between overall editing levels at relapse (AEI at TP2) and editing-to-mutation matches led us to test whether changes in AEI at different time points could be correlated to mutation levels. Indeed, we found that our cohort could be split into two subgroups: in the first, AEI decreases in TP2 (Fold Change <1); in the second, it increases (Fig.3B). We then asked if these two groups also differed, with regard to ADAR1-correlated mutations. Statistical analysis showed that the T-derived mutations within the TP1 editing-defined windows (±20bp) are enriched for patients whose AEI *decreases* in TP2 (Fisher’s exact test: p<10^−3^, Odds Ratio>1, Fig. 3C, Suppl. Tables S1-2). When the editing defined windows were extended by ±100bp, the same analysis showed that the T-derived mutations within the defined windows continue to be significantly enriched for the patients whose AEI decreases (p-val<2.2×10^−16^, Odds Ratio >1) compared those with AEI increase (p=10^−2^, Odds Ratio >1) in TP2 (Fig. 3C, Suppl. Tables S1-2). A few examples of such T-derived unique mutations upon relapse found within the editing-defined windows are provided in Fig. 3D). One broad interpretation of these findings is the possibility that, upon the introduction of an ADAR-catalyzed driver mutation, a tumor loses its need for the presumed benefit RNA editing provides to tumor evolution. Mechanistically, the driver mutation may functionally disrupt ADAR1 targeting on the transcript (e.g. by altering its structure, or the occupancy of RNA binding proteins), thus introducing an additional variable to disease progression. Accordingly, ADAR1 may facilitate tumor adaptation either through the introduction of mutations or through an increase of ADAR editing activity on the mRNA.

**Figure 3.**
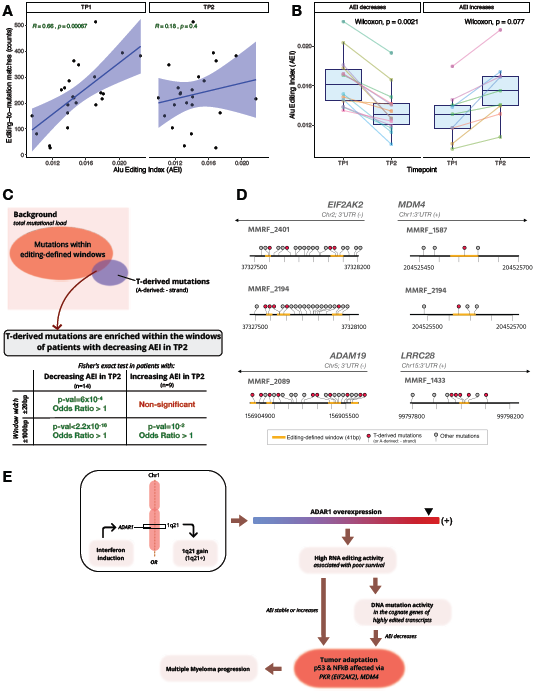
ADAR1 is contributing to Multiple Myeloma progression either through mutation fixation or increased RNA editing activity. (A) Editing-to-mutation matches correlate with the the Alu Editing Index (AEI) of TP1 (p=0.00067), but not with the AEI of TP2. (B) Patients are grouped by whether their AEI decreases or increases in TP2 when compared to TP1 and the first group shows statistically significant lower AEI in TP2 (p=0.0021). (C) T-derived unique mutations in TP2 that fall within the ±20bp windows defined by editing events in TP1, are significantly enriched in patients whose AEI decreases in TP2 (Fisher’s exact test for count data: p=0.00067, Odds Ratio >1) and not for the patients whose AEI increases in TP2. Windows are extended by ±100bp and T-derived unique mutations are more enriched in the windows of the patients whose AEI decreases (p-val<2.2×10^−16^, Odds Ratio >1) than those whose AEI increases (p=10^−1^, Odds Ratio >1) in TP2. (D) Schematic representation (“Lolliplots”) of the unique mutations in TP2 within the given regions of the *EIF2AK2, MDM4, ADAM19* and *LRRC28* genes. Colored with red the T-derived and A-derived mutations for the positively (+) and negatively (-) oriented genes in the genome. The ±20bp windows defined by editing events in TP1 are the blue-colored tracks. The majority of T-derived mutations (or A-derived depending on the gene orientation), are in within or near the conservatively defined windows. (E) The proposed model of ADAR1 as an RNA editor and a DNA mutator in MM: ADAR1 is overexpressed in MM either via 1q21 amplification or through IFN induction. High RNA editing activity in MM may or may not result in DNA mutations in cognate genes with simultaneous decrease in editing activity. Either through increasing/stable editing activity or through mutation fixation and decreasing editing activity the MM tumors readapt; key components, such as PKR or MDM4 that interact with p53 or NFkB, that play a prominent role in MM progression, are regulated.

### Endogenous ADAR1 can be co-opted to mutate DNA

Thus far we have used patient derived tumour data from pre- and post-treatment/relapse timepoints, to correlate ADAR1 editing activity on a transcript with specific types of DNA mutation on the cognate locus. To conclusively demonstrate that ADAR1 is capable of mutating DNA, we asked whether targeted recruitment of ADAR1 to a locus could result in DNA alterations.

Targeted recruitment of ADARs to specific mRNAs can be accomplished with a variety of recently published methodologies (Montiel-González, Vallecillo-Viejo and Rosenthal, 2016; Cox et al., 2017; Vogel et al., 2018): we opted for a simple approach where long, ADAR-recruiting RNAs (or arRNAs) can be designed to engage endogenous editing enzymes and target them to specific sites within mRNAs of interest (Qu et al., 2019). We then hypothesized that this same methodology could be used to recruit ADAR1 to a gene of interest, where ADAR1 would make a specific type of mutation (Fig. 2A). As a straightforward readout, we decided to target the expressed V region gene of the Ramos Burkitt’s lymphoma B cell line, from where the Activation Induced Cytidine Deaminase (Aicda) was deleted. AID protein (the product of the Aicda gene) is responsible for somatic hypermutation (SHM) of antibody genes (Muramatsu et al., 2000). Ramos B cells hypermutate constitutively the V(D)J segment of their productively recombined antibody gene (Sale et al. 1998, Cook et al., 2007 and Fig. 4A: AID WT), and loss of AID leads to an inability of these cells to hypermutate – consequently, AID-/- cells are unable to lose surface IgM expression (which is a measure of hypermutation - Cook et al., 2007 and Fig. 4A: AID-/-). Finally, transfection with vectors that express AID cDNA can reconstitute the IgM-loss phenotype (Fig. 4A, AID-/- mAID and Zhang et al., 2001; Al-Qaisi, Su and Roffler, 2018).

**Figure 4.**
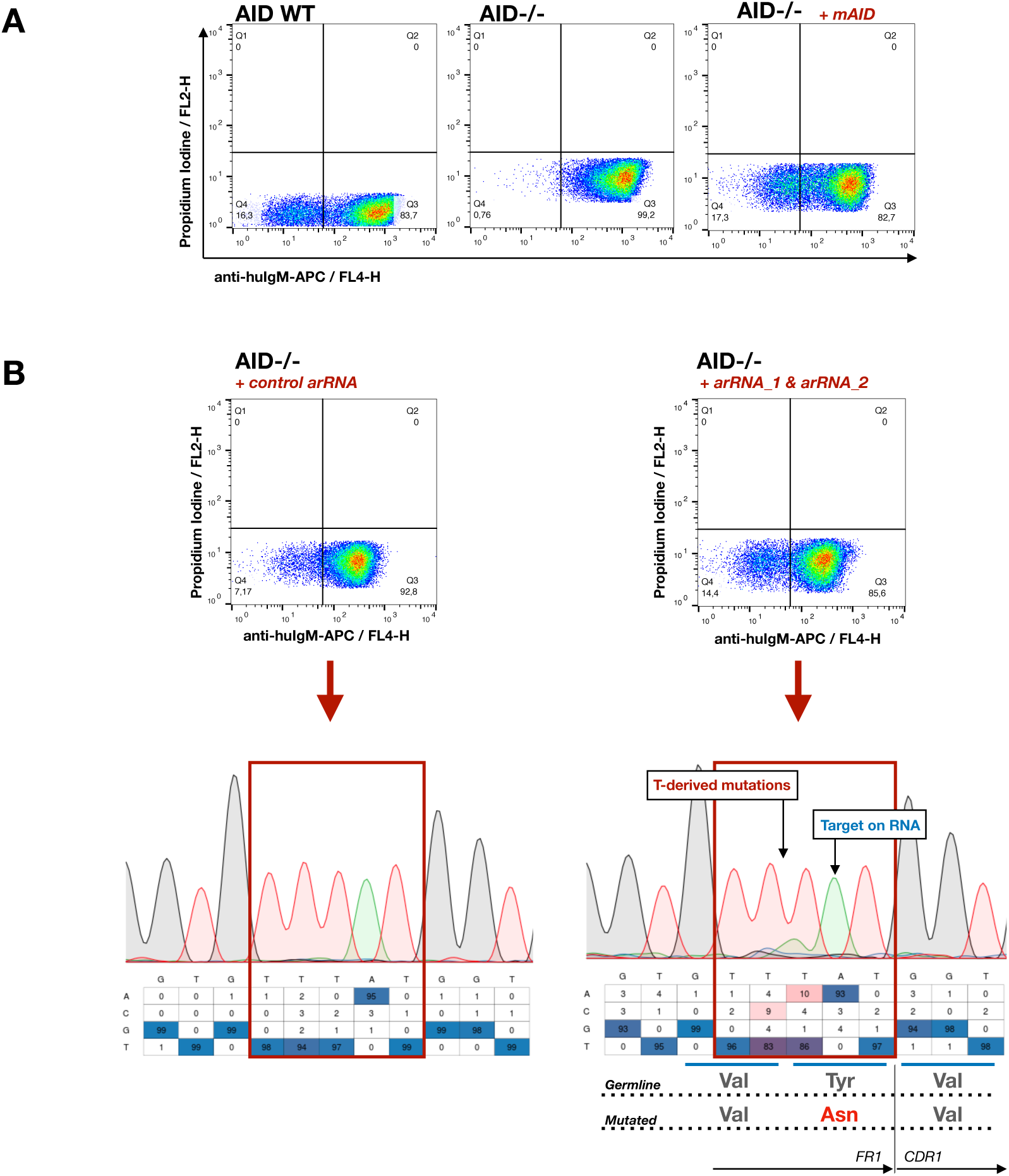
Endogenous ADAR1 is recruited to the variable region of the expressed antibody gene in Ramos cells, where it mutates gDNA. (A) Ramos AID wild-type (WT) cultures comprise both an IgM+ and IgM-population (Q4, 16.3%). In AID-/- Ramos cells, the IgM-population is completely absent, regardless of the time the cells have been in culture (Q4, <1%). IgM-loss can be reconstituted in AID-/- cells upon transient expression of vectors carrying the cDNA of mouse AID (mAID) (Q4, 17.3%). (B) AID-/- cells transfected with two ADAR-recruiting RNAs (arRNA_1 & arRNA_2) show increased IgM-population versus cells transfected with an arRNA targeting thymidine kinase (which does not exist in the Ramos genome - control arRNA). T-derived mutations are reported in gDNA amplicons (derived from bulk culture), in the vicinity of where ADAR1 was targeted on the RNA of the Ramos V-region. T-to-A mutation adjacent to the A-target, is in the first position of the last codon of the Framework Region 1 (FR1; according to IMGT), which leads to amino-acid change from tyrosine to asparagine. The gating strategy of the FACS plots is shown in Fig. S5.

To ask whether endogenous ADAR1 can be recruited to the Ramos V region gene segment we designed a series of 71nt long arRNAs to target a specific adenosine (A) on the transcribed strand. Plasmids expressing these arRNAs were transfected into cells using electroporation together with plasmids expressing GFP, and green cells were FACS-sorted and allowed to recover and expand for at least 2 weeks before they were queried for surface IgM loss. We found that a combination of two arRNAs could indeed produce IgM loss when transfected into an AID-/- Ramos B cell line (Fig. 4B; compared to lines transfected with unrelated guides). Sequence analyses of DNA amplicons from the targeted region (amplified from bulk DNA) show the position of the detected T-derived mutation vs the location of the targeted adenosine (Fig. 4B, chromatographs). On the protein level, the T-to-A mutation on the DNA amplicon, right before the target-position on the RNA, is the start of the last codon of the Framework Region 1 (FR1; according to IMGT database - www.imgt.org) and leads to an asparagine-to-tyrosine change on the protein level. This change may explain the IgM loss shown in this experiment, since it occurs at the junction between FR1 and the start of the Complementarity Determining Region 1 (CDR1), a region important for antibody stability. Mutations in the same set of codons have been previously shown to contribute to IgM loss (Sale et al., 1998; Rada, Jarvis and Milstein, 2002). Notably, guide-expressing plasmids were lost from the culture (through dilution) by the time of this analysis, and consequently, RNA editing could not be detected at the level of cDNA (data not shown), consistent with the idea that ADAR-guided IgM loss (through targeting DNA) occurred early and was clonally propagated.

While further optimization of targeting protocols is warranted and ongoing, these experiments demonstrate that endogenous ADAR1 is capable of targeting genomic DNA, with reasonable efficiency.

## DISCUSSION

Multiple myeloma aggressiveness and progression has long been associated with a specific chromosomal translocation (1q21). This is also the location of the gene *Adar1*, which encodes a key RNA editing enzyme that is of emerging relevance to disease progression. Consequently, a number of laboratories have studied the role of ADAR1 mediated RNA editing in MM, establishing editing as a key prognostic indicator (Lazzari et al., 2017; Teoh et al., 2018 and **Fig 1**). Here we have focused specifically on the role of ADAR1 mediated deamination in the context of MM disease *progression*. For this, we curated a group of patients for whom we had access to good quality RNAseq and WES data during two timepoints (pre and post relapse or treatment). A comparative analysis of editing in these matched datasets defined new targets of ADAR1 editing (e.g. the transcripts that encode PKR and MDM4) with potential functional consequences for disease progression.

More importantly, our analyses of these matched datasets revealed a concurrence between editing and the acquisition of new mutations at genes whose cognate transcripts are edited, and at specific predetermined locations surrounding the site-to-be-edited. Such locations are defined by the edited adenosine within a transient RNA:DNA hybrid (an R-loop) with an average size range of ∼80bp-300bp – (Chen et al., 2019; Malig et al., 2019; Stolz et al., 2019) direct DNA A-to-I deamination would then be centered around the deoxyadenosine whose ribo-adenosine counterpart is edited in mRNA (e.g. an adenosine in the 3’UTR), converting it to deoxy-inosine (hypoxanthine). This process would contribute to T-to-C (when read from the non-transcribed (+) strand) or generally T-derived mutations if DNA repair has occurred (Fig. 3). This correlation, paved the way for our experimental validation; guide-based re-targeting of endogenous RNA editors to specific genomic locations (Qu et al., 2019) has the capacity to generate specific mutations at predetermined locations. This initial experimental validation (Fig. 4), in which T-derived mutations in gDNA are found in the vicinity of the targeted adenosine on the transcript, is experimental proof that endogenous ADAR1 is capable of being redirected to DNA (possibly in the context of a short, guide-induced R-loop) to generate DNA mutation. Further optimization is required to generalize this approach for the purposes of introducing targeted DNA mutation using endogenous editing enzymes together with exogenously provided guide RNAs.

Overall, our data suggest that the RNA editing enzyme ADAR1 is likely to function as a DNA mutator during multiple myeloma (MM) progression (Fig. 3E). Our findings are coherent with recent analyses suggesting potential contributions of ADAR1 mediated T-to-C mutation at variable regions of antibody genes, through direct DNA A-to-I deamination at transcription bubbles (Steele and Lindley, 2017) and with population genomics approaches that associate signatures of selection in both coding and non-coding regions of the genome with A-to-I editing (Popitsch et al., 2017; Breen et al., 2019). On a broader spectrum, concurrence of RNA and DNA alterations have been recently reported in several cancers (PCAWG Transcriptome Core Group, et al., 2020), but with no clear suggestions about their sources. Our results agree with RNA/DNA modifications concurrence and, moreover, link them directly under the suggested dual role of ADAR1 *in vivo*.

The ADAR1-attributed mutations we identified in TP2 that matched editing events defined locations in TP1 occurred at a reasonably high frequency (5% of the tumor, on average). However, they remain subclonal, since in a few cases TP2 was a timepoint in which the patients were sequenced for a post-treatment check. It is possible that at later time-points these subclonal mutations would confer a selection advantage (especially as they are present in genes key to MM progression – Fig. 3). At the same time, while it is known that APOBECs (such as APOBEC1 and multiple APOBEC3 family members) are the source of DNA mutations in cancer (estimated to occur at the rate of ∼1/10,000bp per generation (Saraconi et al., 2014), our data imply that these mutations are not randomly distributed in the genome, but are in fact centered on the genes whose transcripts are targets of the RNA editing activity of the relevant APOBEC enzyme.

Overall, our data suggest that the dual role of RNA editor and DNA mutator might be shared by many deaminases, so that DNA mutation might be the result of collateral damage on the genome by an editing enzyme, whose primary job is to re-code the cognate transcript toward specific functional outcomes.

## METHODS

### RNA-seq and WES data processing

RNA-seq and WES data from 590 patients were retrieved from the CoMMpass MMRF study (dbGaP accession number phs000748; http://www.ncbi.nlm.nih.gov/gap). RNA-seq and WES data were mapped against the human reference genome GRCh37 and all annotation and gene models were based on Ensembl version 74. RNA-seq data was mapped with GSNAP (v. 2017-06-20; PID: 15728110) using the default parameters, duplicate reads were marked with Picard (v 1.93; http://broadinstitute.github.io/picard/), and aligned data were sorted and indexed with Samtools (v. 0.1.19; Li et al., 2009). RNA-seq unmapped reads were realigned to include hyperedited reads as preciously described (Porath, Carmi and Levanon, 2014). WES data was processed as recommended by the ‘GATK Best Practices’ as previously described (Laganà et al., 2018).

### RNA editing analyses and WES mutation calling

The REDItools suite was used to detect RNA editing sites (Picardi and Pesole, 2013; Picardi et al., 2015). In particular, the REDItoolDenovo.py python script was employed to detect Single Nucleotide Variations (SNVs) when compared to reference genome. Only well-covered (≥10 reads) and reported as statistically significant (p-value≤0.05) SNVs in concordant read-pairs were considered for downstream analyses. RNA editing events were defined after genomic positions with mutations called from WES data were subtracted from the SNV genomic coordinates. Mutations were called from WES data employing Strelka2 (Kim et al., 2018) and Vardict (Lai et al., 2016). Mutations at any frequency, but well-covered (≥10 reads), were considered for proper exclusion of any DNA signal resembled on the RNA.

For the editing-to-mutation analysis, we focused on a subset of 23 patients (Fig. 2B) that had a complete set of RNA-seq and WES for two successive timepoints (TP1 and TP2). Downstream analysis was performed so as to match RNA editing events for each patient in the TP1 to unique mutations in TP2. The matching candidates were validated with the tool *bam-readcount* (https://github.com/genome/bam-readcount) and annotated with Oncotator v.1.9.9.0 (Ramos et al., 2015).

### AEI, expression/pathway analyses

Pathway enrichment analysis was performed with SLAPenrich (Iorio et al., 2018). Alu Editing Index (AEI) was calculated from RNAseq data using REDItools as described above for detection of RNA editing sites and in-house python scripts implementing a previously described procedure (Bazak, Levanon and Eisenberg, 2014; Paz-Yaacov et al., 2015; Roth, Levanon and Eisenberg, 2019).

Copy number estimates were identified from the long-insert WGS results using existing TGen tools, as previously described (Laganà et al., 2017).

### IgM loss through DNA mutation recruiting the endogenous ADAR1

Ramos AID-/- cells were transfected with plasmids containing arRNAs (pENTR promoter) and pMax (GFP, Amaxa, Lonza) at ratio of 1:10, with total 2ug plasmid DNA. 48h upon transfection cells were sorted for GFP with BD FACSAria II and after two weeks, fraction of the cell population was stained for Propidium Iodine (PI) to observe cell viability and APC-conjugated, Goat, Anti-Human IgM (Jackson Immunoresearch) to observe IgM loss For flow cytometry Calibur (BD) was used. The other fraction of the cell population underwent simultaneous gDNA/RNA extraction with AllPrep DNA/RNA Mini Kit (Qiagen). The extracted RNA was treated with TURBO DNAse (ThermoFisher Scientific). The gDNA amplicons from the V-region were amplified with the Q5 High-Fidelity DNA polymerase (New England Biolabs) protocol, while the cDNA amplicons were made with OneStep RT-PCR kit (QIAGEN). Manufacturer protocols were followed for all experiments. Primers for the cDNA amplicons were from Tiller et al., 2008, while gDNA from Sale et al., 1998. the Sanger Sequencing chromatograms were all quantified SNVs with MultiEditR (Kluesner et al., 2019).

## AUTHORS CONTRIBUTIONS & ACKNOWLEDGMENTS

The work described herein was conceived by RT, AL, SP and FNP, based on data initially produced in the context of a different project spearheaded by RP, FNP in collaboration with Prof Qiang Pan-Hammarström at Karolinska Institute, Sweden, whom the authors thank. RT, AL performed all data analyses. All authors contributed to writing the manuscript. SP and FNP wish to thank Dr Rita Shakhnovich, who suggested the collaboration between them. The authors also wish to thank Dr. Jeroen Guikema, UMC Amsterdam, The Netherlands for the gift of AID-/- cell lines and the Flow Cytometry facility of DKFZ, Heidelberg, Germany. Finally, authors also wish to thank Prof Silvo Conticello and Dr Salvatore di Giorgio at Tuscan Tumor Institure, Florence Italy, for their comments on the manuscript.

