## Supplementary Material for "ADAR1 can drive Multiple Myeloma progression by acting both as an RNA editor of specific transcripts and as a DNA mutator of their cognate genes"

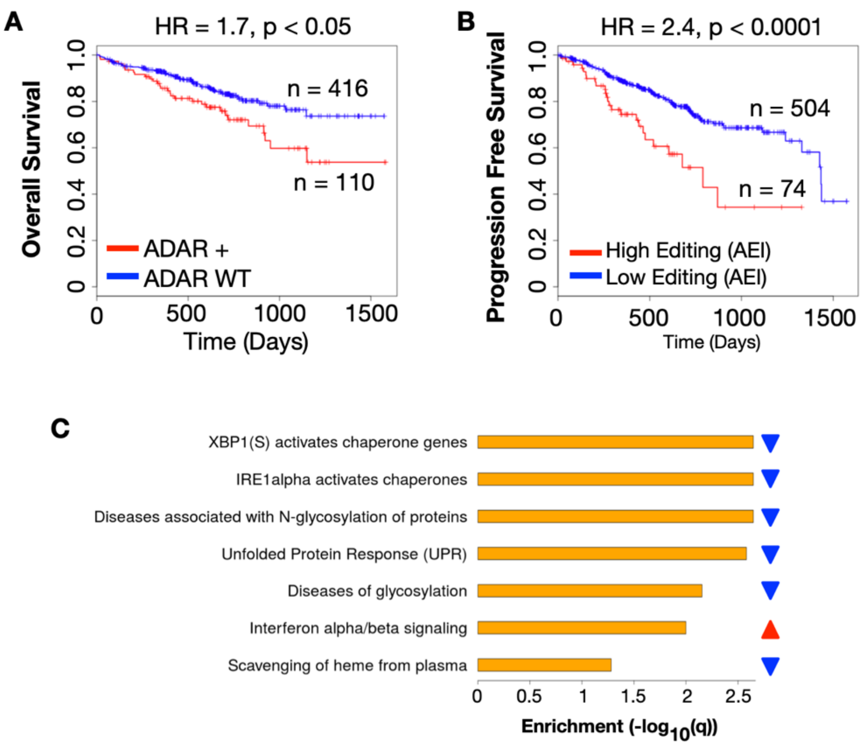


**Figure S1. ADAR1 copy number and editing activity is associated with poor prognosis in Multiple Myeloma, affecting transcripts of genes involved in protein response and processing.** (A) Overall survival of patients with ADAR1 copy number (CN) gain (>2) (ADAR+, n=110) and ADAR1 Wild Type patients (ADARwt, n=416), (B) Progression-free survival of patients with high AEI (Alu Editing Index; n=74) and low AEI (n=504), (C) pathway enrichment analysis in the patients with high AEI shows up-regulation of interferon alpha/beta signaling and down-regulation of pathways involved in protein response (UPR, XBP1(S) and IRE1alpha activate chaperone genes) and glycosylation (N-glycosylation and glycosylation of proteins associated diseases).

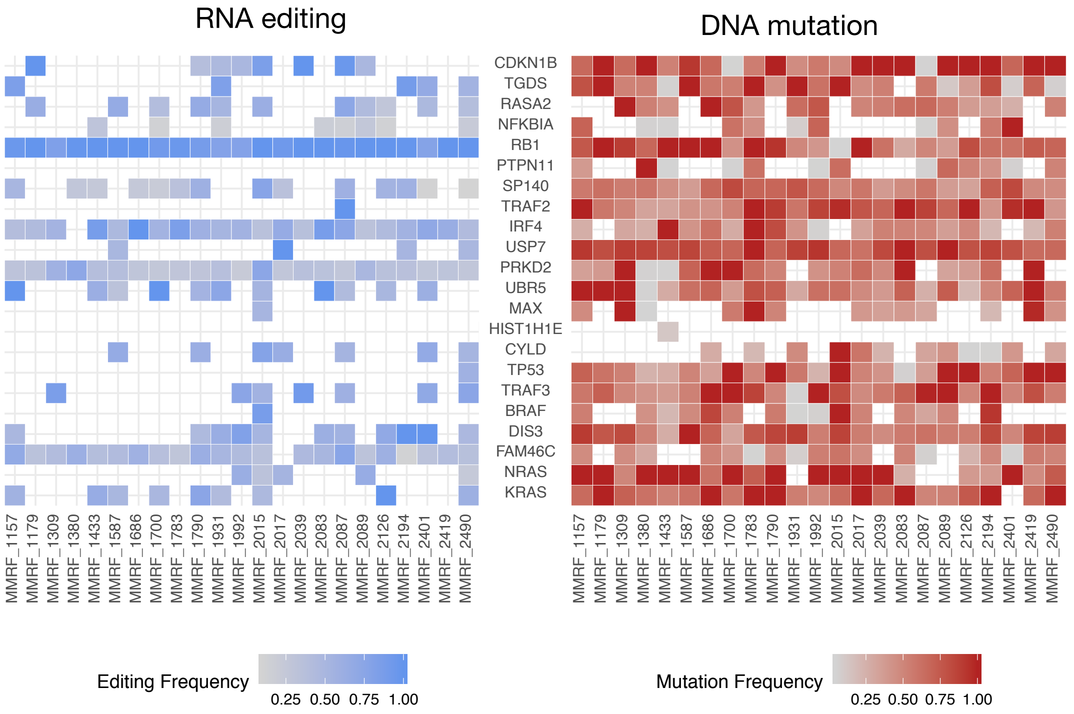


**Figure S2: RNA editing on transcripts and mutations of their cognate genes in known Multiple Myeloma drivers** (Walker et al., 2018) **in TP1.** Heatmaps tally up per patient the average variation frequency between different genomic positions across the exons of each transcript or gene. Mutation heatmap visualizes all substitutions detected (cut-off 5% variation frequency and only good quality reads were considered), while RNA editing visualizes A-to-I editing by ADAR1.

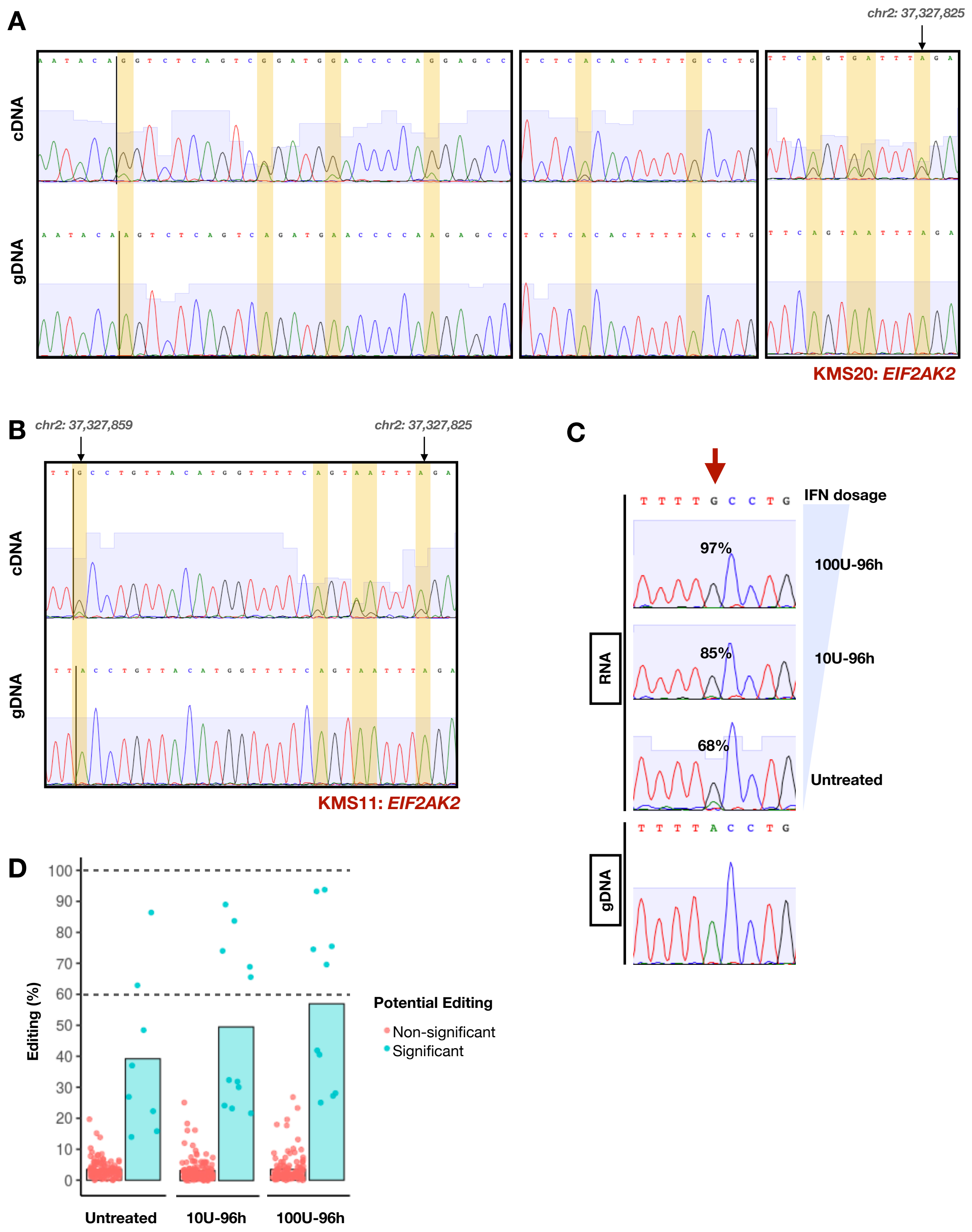


**Figure S3: Validation of RNA editing in EIF2AK2** in KMS20 (A) and KMS11 (B) MM cell lines by comparing RNA and DNA, simultaneously extracted by untreated cells from the cell culture. Sites detected in cell lines greatly overlap with genomic coordinates detected in the patient data from the MMRF study. When KMS11 cells are treated with 10U and 100U of IFNα+γ for 96h, editing efficiency drastically increases. Panel (C) shows such a position in the 3'UTR of EIF2AK2, where Editing % increases from 68% in untreated cells to 97% in 100U IFN-treated cells for 96h. (D) Editing % in aggregate throughout the 3'UTR region sequenced increases with IFN-treatment of the cells. Sanger sequencing chromatograms from the amplicons were quantified with MultiEditR (Kluesner et al., 2019). Bar plots indicate the mean of Editing (%) of the significant sites (default paremeters and test; Kluesner et al., 2019).

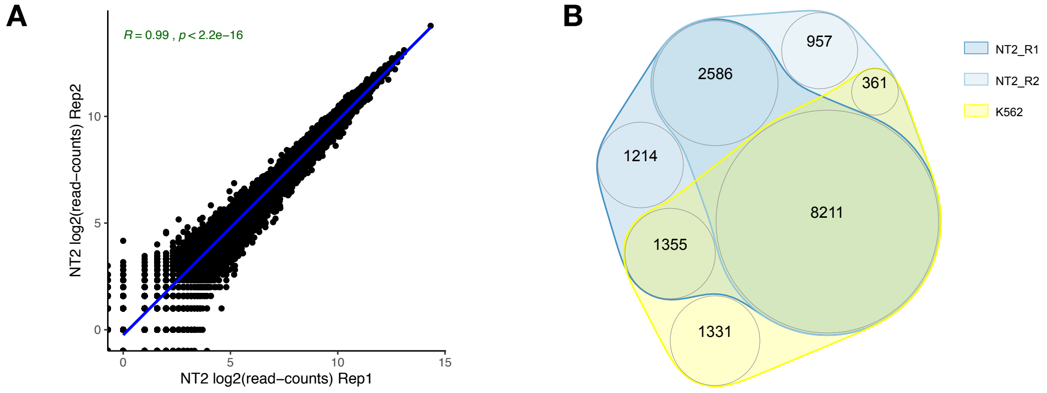


**C**

**How many of the editing-to-mutation matches fall in the DRIP-seq R-loops?**

- **80.5% of the matches fall into the Union of NT2_R1, NT2_R2 & K562 (Union = 167902 R loops, formed by 16015 transcripts to the gene annotation).**
- **66.2% of the matches fall into the overlap of NT2_R1, NT2_R2 & K562 (Overlap = 139811 Rloops formed by 8211 transcripts).**

**Figure S4: Mapping of R-loops (publicly available DRIP-seq data: GSE70189) on the editing-to-mutation candidates.** (A) The expression profiles of replicates of the NT2 cell line correlate significantly ($R=0.99$, $p<2.2x10^{-16}$). (B) There's a great overlap of the transcripts (gene-level annotation) that reportedly form R-loops between the NT2 cell-line replicates and K562 (no expression data available). About 72% of K562 R-loop-forming transcripts overlap with NT2 transcripts that form R-loops. (C) Mapping of the R-loops on the editing-to-mutation candidates, indicates that 80.5% of them falls into the union of the different cell-line sets tested and 66.2% into the absolute overlap of the cell-line sets tested. Datasets come from GEO with accession number GSE70189 (Sanz et al., 2016; Sanz and Chédin, 2019).

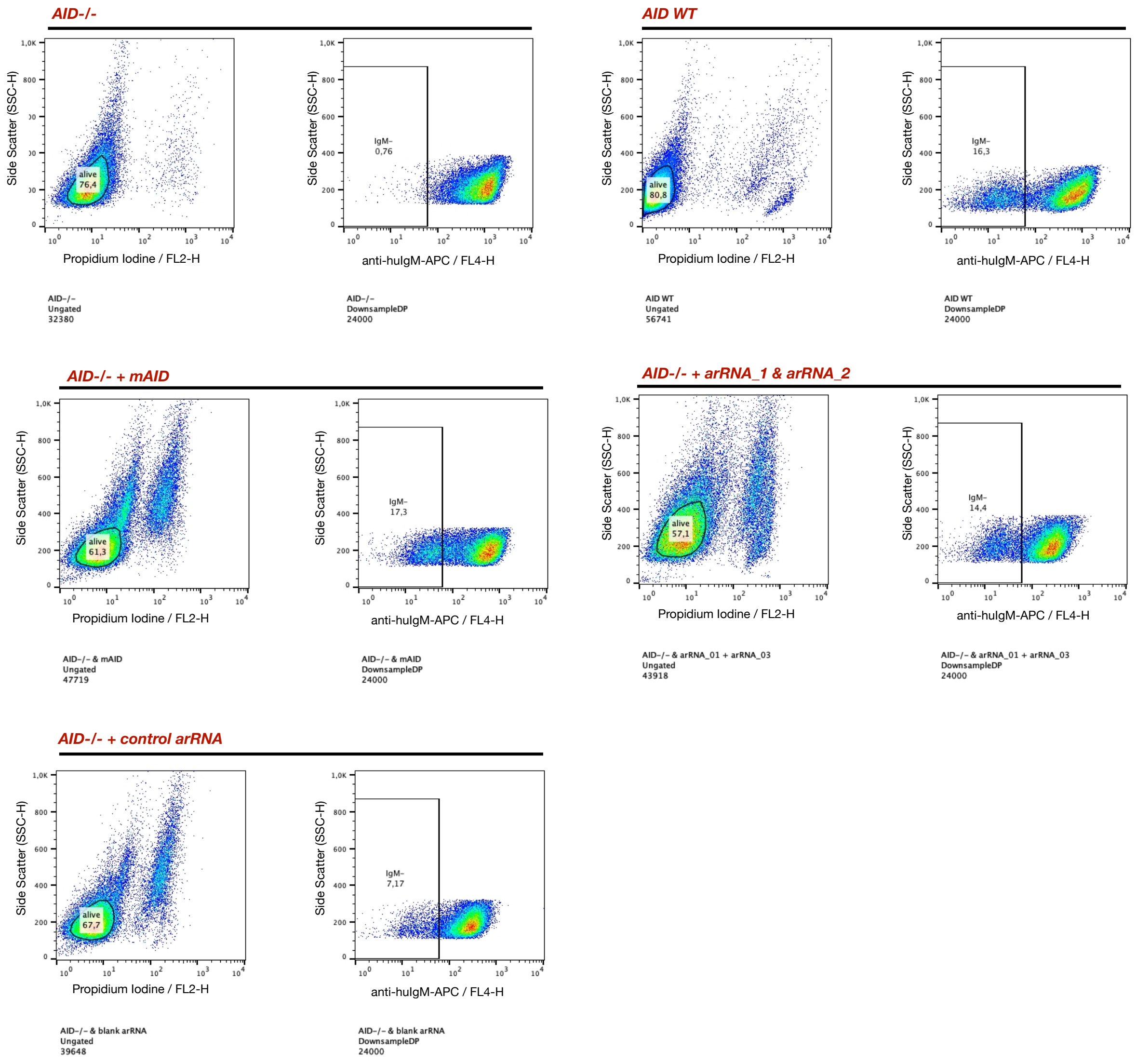


**Figure S5: Gating strategy for flow cytometry data.** Cell populations were stained with propidium iodide (PI) – absorbing at FL2 - and with APC-fluorescent IgM-specific antibody - absorbing at FL4. Initially, the background of dead-cell population was eliminated by selecting the PI-negative population. Then, the different populations were down-sampled to an equal number (24000) and gated into FL4 for sorting out the IgM-positive from the IgM-negative (IgM-) populations. Related information with figures 4 and S4.

**Table S1: Contingency tables summary for the unique in TP2 T-derived mutations significance in ±20bp and ±100bp editing-defined windows in MM.** Related information with figure 3C.

| | | AEI TP2 < AEI TP1 | | AEI TP2 > AEI TP1 | |
|---|---|---|---|---|---|
| | | Out of window | In the window | Out of Window | In the window |
| ±20bp | No T-derived | 42184389 | 17495 | 24097508 | 8021 |
| | T-derived | 9393398 | 4133 | 5381380 | 1744 |
| ±100bp | No T-derived | 42134647 | 67237 | 24074484 | 31045 |
| | T-derived | 9381203 | 16328 | 5375954 | 7170 |

**Table S2: Fisher's exact test for count data results and parameters for contingency tables in Table S1.** Related information with figure 3C and Table S1.

| | | | **95% confidence interval** | |
|---|---|---|---|---|
| **Test group** | **p-value** | **Odds Ratio** | **Min value** | **Max value** |
| **±20bp / AEI TP2 < AEI TP1** | 0.0006709 | 1.060925 | 1.025283 | 1.097567 |
| **±20bp / AEI TP2 > AEI TP1** | 0.3194 | 0.9736315 | 0.9239634 | 1.0255253 |
| **±100bp / AEI TP2 < AEI TP1** | < 2.2e-16 | 1.09072 | 1.072137 | 1.109533 |
| **±100bp / AEI TP2 > AEI TP1** | 0.01035 | 1.034262 | 1.007855 | 1.061216 |

Walker, B., Mavrommatis, K., Wardell, C., Ashby, T., Bauer, M., Davies, F., Rosenthal, A., Wang, H., Qu, P., Hoering, A., Samur, M., Towfic, F., Ortiz, M., Flynt, E., Yu, Z., Yang, Z., Rozelle, D., Obenauer, J., Trotter, M., Auclair, D., Keats, J., Bolli, N., Fulciniti, M., Szalat, R., Moreau, P., Durie, B., Stewart, A., Goldschmidt, H., Raab, M., Einsele, H., Sonneveld, P., San

Miguel, J., Lonial, S., Jackson, G., Anderson, K., Avet-Loiseau, H., Munshi, N., Thakurta, A. and Morgan, G. (2018). Identification of novel mutational drivers reveals oncogene dependencies in multiple myeloma. Blood, 132(6), pp.587-597.
